# Seasonal fluctuations in fitness result in severe reductions in effective population size

**DOI:** 10.64898/2026.03.30.715388

**Authors:** Olivia L. Johnson, Raymond Tobler, Joshua M. Schmidt, Christian D. Huber

## Abstract

Genetic evidence for fluctuating selection has begun to accumulate for different species over the past few decades, especially for the *Drosophila* genus where studies have reported hundreds of loci undergoing putatively adaptive oscillations across successive seasons. However, most theoretical and simulation studies of fluctuating selection have relied on abstract or weakly parameterized models, making it difficult to assess their relevance for natural populations. In this study, we simulate multilocus seasonally fluctuating selection under a recently developed model and examine its effect on the variance effective population size (*N_e_*) at a genome-wide scale. By recapitulating genomic, demographic, and evolutionary parameters from natural *Drosophila* populations in our simulations, we were able to reproduce allele frequency oscillations reported in recent studies and show that these lead to ∼50% genome-wide reductions in *N_e_*. We also demonstrate that *N_e_* reductions are well predicted by the maximum frequency amplitude among all adaptively fluctuating loci, and that the frequency amplitudes are largely determined by the number of adaptively fluctuating loci and the strength of their epistatic interactions. Our results demonstrate that fluctuating selection can substantially reduce effective population size and underscore the importance of temporally variable selection in shaping genome-wide patterns of variation beyond classical models.

**Article Summary:** Genetic studies of fluctuating selection in natural populations have grown steadily over the past decade, with reports suggesting that hundreds of loci undergo adaptive oscillations over seasonal timescales in cosmopolitan *Drosophila* populations. By simulating seasonally fluctuating selection under a recently developed model and ecological scenarios informed by published studies, the authors show that this mode of selection can reduce effective population size by ∼50%, with the magnitude of the reduction correlated with the locus exhibiting the largest allele frequency fluctuations. These findings highlight fluctuating selection as an important factor shaping genome-wide patterns of genetic variation and effective population size.

## Introduction

Effective population size (*N_e_*) is a central parameter in population genetics, governing the strength of genetic drift and shaping patterns of genetic variation. Although *N_e_* is often interpreted in terms of demographic processes, it can also be strongly influenced by selection through its effects on the distribution of reproductive success among individuals. Different formulations of *N_e_* capture these dynamics from distinct perspectives, including variance *N_e_*, which reflects fluctuations in reproductive success across generations, and coalescent *N_e_*, which is inferred from levels of genetic diversity (Crow 1954; Sjödin et al. 2005; Wakeley & Sargsyan 2009; Wang et al. 2016; Waples 2022).

Selection is known to systematically alter these measures. Positive selection and purifying selection can reduce *N_e_* by increasing variance in reproductive success and by depleting linked neutral diversity through selective sweeps and background selection (Charlesworth 2012; Comeron 2014; Corbett-Detig et al. 2015; Comeron 2017; Buffalo 2021; Charlesworth & Jensen 2022; Achaz & Schertzer 2023). In contrast, balancing selection can maintain genetic variation and thereby increase estimates of *N_e_* (Charlesworth 2006; Rodríguez de Cara et al. 2023).

One selective process that has received increased attention recently but remains relatively unexplored in relation to effective population size is fluctuating selection – i.e., adaptive scenarios where selection changes in direction or intensity across time (Dobzhansky 1947; Haldane & Jayakar 1963; Gillespie 1991; Johnson et al. 2023). A number of theoretical and empirical studies have recently demonstrated that fluctuating selection can have wide-ranging effects on levels of population genetic diversity (Barton 2000; Taylor 2013; Huang et al. 2014; Wittmann et al. 2023). In particular, a recent theoretical study reported a 30% reduction in genome-wide diversity when seasonal fluctuating selection acted upon a single site, with these impacts also extending across unlinked neutral sites on different chromosomes (Wittmann et al. 2023). This remarkable genome-wide reduction in diversity was caused by the onset of serial reproductive bottlenecks occurring at the start of each season, where the least fit individuals from the previous season exhibited a significant reproductive advantage in the present season, resulting in greatly increased variability in offspring number between individuals (Wittmann et al. 2023).

These results indicate that species susceptible to temporally fluctuating selection pressures may also be expected to experience a severe decrement in *N_e_.* Moreover, while our understanding of the scope and impact of fluctuating selection in natural populations is currently limited to studies conducted on a handful of species (Kelly 2022; Pfenninger & Foucault 2022; Lynch et al. 2024; Johnson et al. 2023), research on natural and experimental outdoor populations of *Drosophila* suggest that hundreds to potentially thousands of fluctuating loci are under seasonal selection in these cosmopolitan taxa (Bergland et al. 2014; Behrman et al. 2018; Machado et al. 2021; Rudman et al. 2022; Bitter et al. 2024; Nunez et al. 2024). However, the impact of this pervasive fluctuating selection signal on *N_e_* remains unknown, and our ignorance further extends to how factors like the number of fluctuating loci, selection strength, and census population size combine to mediate impacts on *N_e_*. In the present study, we employ forward simulations that draw upon a recently published multi-locus model of seasonal adaptation (Wittmann et al. 2017) and parameter ranges informed by empirical studies of *D. melanogaster* (Bergland et al. 2014; Machado et al. 2021; Rudman et al. 2022; Bitter et al. 2024) to provide new insights into the impact of fluctuating selection on instantaneous variance *N_e_* under plausible evolutionary scenarios.

## Results

### Simulating adaptive seasonally fluctuating loci

To assess the impact of fluctuating selection on *N_e_*, we used the forward population genetic simulator *SLiM* (v 4.0.1; (Haller & Messer 2023)) to replicate multi-locus adaptation across two alternating seasonal environments (i.e., summer and winter; s/w) under the Segregation Lift (SL) model developed by Wittman and colleagues (Wittmann et al. 2017). The SL model comprises a dynamic fitness landscape where the seasonal scores for each individual (i.e. *z_s_* for summer and *z_w_* for winter; see equation 1) are obtained by summing the contributions of all seasonally adaptive loci (each locus-specific contribution, *C_l_*, being the product of a seasonal dominance coefficient (*d_s/w_*) and effect size (Δ*_s/w_*); Table 1; see Methods). This score is converted to fitness, *w*(*z*), via a power transformation (scale = *y;* equation 2) that allows for non-additive contributions across seasonal loci (i.e. epistasis; also see Methods for further discussion).

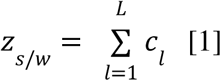

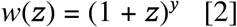

For our initial simulations, we focused on quantifying the dynamics of allele frequency fluctuations of seasonally adaptive loci (henceforth seasonal loci) under a simple demography where population size remained constant at one million individuals across 36,000 generations and each season had 10 generations. Note that we were able to improve the computational efficiency of our simulations while still obtaining representative results by downscaling each of these parameters by a factor of two (see Methods) but refer to the unscaled parameter values throughout for convenience.

Initially, we sought to identify the impact of the number of seasonal loci and epistasis intensity on the magnitude of frequency fluctuations and the number of generations needed to achieve stable oscillations. Accordingly, separate simulations were run using different combinations of both variables – seasonal locus count ranging between 100 to 500 and epistasis intensity (*y*) ranging from 0.5 to 20; ranges based on reports from relevant *D. melanogaster* studies (Bergland et al. 2014; Machado et al. 2021; Rudman et al. 2022; Bitter et al. 2024) – with all loci being introduced in the first generation with each seasonal allele at 50% frequency. Seasonal dominance and effect sizes were randomly drawn from independent distributions (see Methods), facilitating a broad range of seasonal allele frequency fluctuations across all seasonal loci. The fitness function (equation 2) induces diminishing-returns fitness scenarios for all epistasis intensities, as this type of epistasis has prior support from empirical studies (Chou et al. 2011; Khan et al. 2011; Kryazhimskiy et al. 2014; Wittmann et al. 2017) and produces stable fluctuations with plausible properties (Wittmann et al. 2017)(see Methods). After running each parameter combination 20 times, we evaluated trends in allele frequency changes for all seasonal loci sampled over three consecutive seasonal cycles at the commencement, midpoint (i.e., from 18,000 generations), and end of the simulation (i.e., from 36,000 generations).

### Fluctuating allele frequencies depend on the number of segregating seasonal loci and epistasis intensity

Across all simulations, we observe a continuous decline in the number of segregating loci across time (Figure S1), with loci with similar seasonal effect size (i.e. *Δ*_s_ / *Δ*_w_ ≅ 1) being more likely to remain segregating throughout, while those with skewed effects (i.e.| *Δ*_s_ - *Δ*_w_ |>> 1) were more prone to loss (Figure S2), corroborating theoretical results (Wittmann et al. 2017). The rate of seasonal locus loss is strongly dependent on the epistasis intensity, with weakly epistatic loci becoming lost at a steady rate whereas loci experiencing stronger epistasis are lost at higher rates initially, before levelling off toward the end of the simulation.

Despite the continual loss of seasonal loci through time, seasonal allele frequency fluctuations had stabilised by 18,000 generations, with mean allele frequency amplitudes (i.e. seasonal frequency range averaged over successive cycles at each sampling point) ranging between ∼0.4% to 16.5% (see Methods; Figure S3). Notably, the relationship between seasonal locus loss rate and epistasis was maintained after downscaling the population size by an order of magnitude (Figure S4; see Methods), with increased sampling during the early stages of the downscaled simulations revealing that the seasonal allele frequency amplitudes had become stable by 1,200 generations across all parameter combinations (Figure S5). This rapid stabilisation likely extends across all of our simulations, since alleles at all seasonal loci start at 50% frequency from the onset of the simulations, which is the mean value of the oscillating alleles at equilibrium, and the combined results indicate the stable frequency oscillations are maintained even as seasonal loci continue to be lost.

Strikingly, the seasonal frequency amplitudes at the end of the simulations showed a strong inverse correlation with the final number of segregating seasonal loci but a positive relationship with epistasis intensity (Figure 1). After log-transforming each variable, we were able to fit a linear regression model to our simulated data that accurately predicted the mean seasonal frequency from linear combinations of seasonal loci number and epistasis intensity (*r*^2^ = 0.988). Similar fits were obtained for regression models predicting the median (*r*^2^ = 0.963), maximum (*r*^2^ = 0.921), and 90% quartile frequency amplitudes (*r*^2^ = 0.985), indicating that frequency fluctuations in the centre and upper tail of the distribution are highly predictable. Further inspection of the down sampled simulations showed that this linear relationship between the log-transformed oscillating allele frequencies and segregating seasonal loci had converged by 18,000 generations for all epistasis intensities, with convergence being faster for loci with higher epistasis intensities (Figure S6).

**Figure 1.**
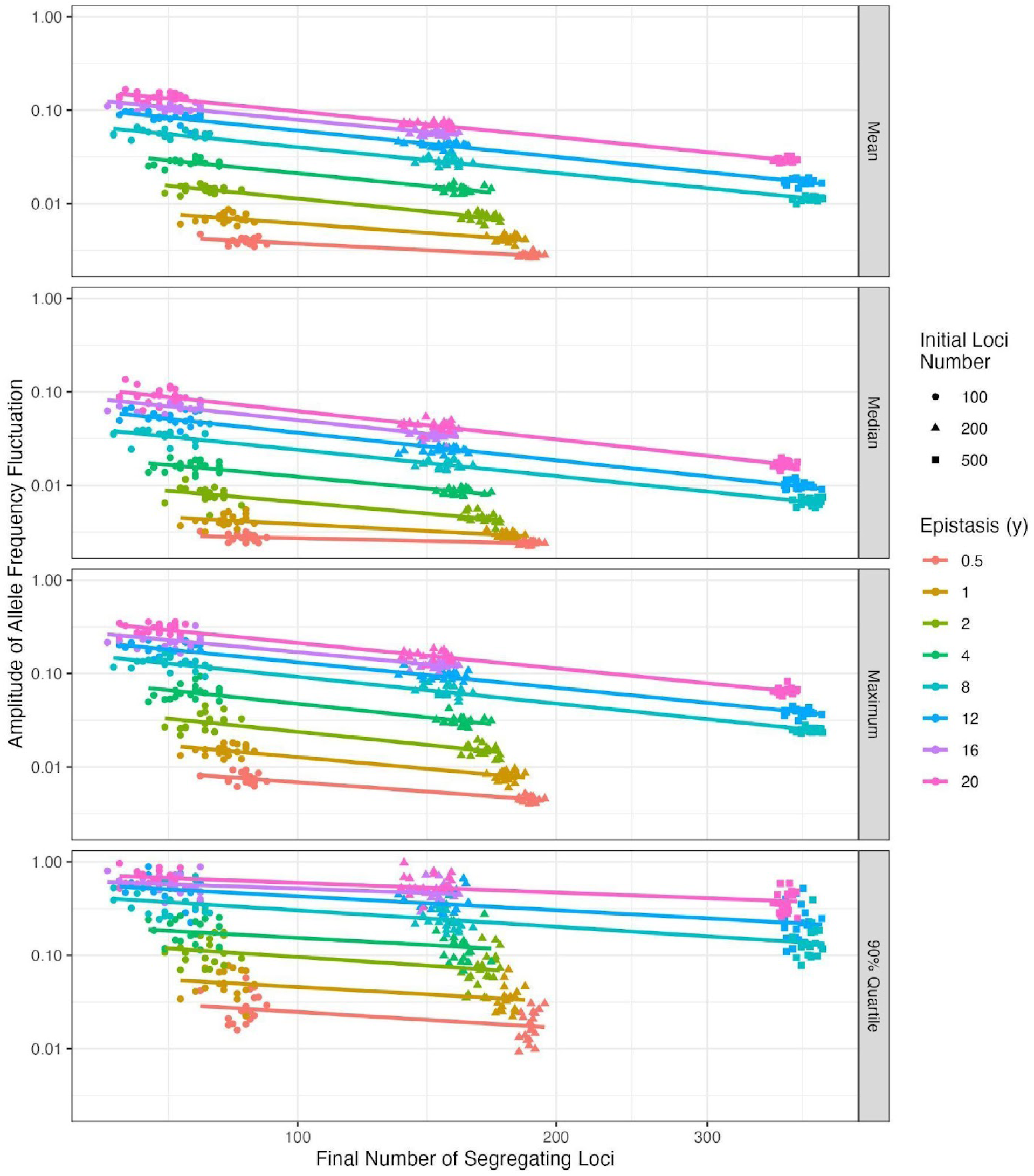
Relationship between number of segregating loci, epistasis, and amplitude of seasonal allele frequency fluctuation. Each panel depicts the relationship between the initial and final number of segregating loci (sampled at 18,000 generations) and the mean, median, maximum, or 90% quartile of allele frequency fluctuations, for different epistasis values. The amplitude of the seasonal fluctuation is shown on the y-axis. The initial loci number is visualised by the shape of the points, with each point representing a single simulated replicate (of which there are 20 for each initial loci number and epistasis value). The colour of the points and lines indicated the epistasis value.

These relationships prompted us to fit regression models to the simulated datasets that predict the expected epistasis intensity as a linear function of the number of segregating seasonal loci and their mean oscillation magnitude (see equations 3 and 4 in Methods). Because the number of segregating loci was also predictable from the number of starting loci (Figure 1), we used the best fitting regression model to estimate the combination of epistasis intensity and initial number of seasonal loci required to replicate the number and amplitude of fluctuating loci reported in natural *D. melanogaster* populations (Table 2; see Methods). The estimated number of seasonal loci and interaction intensities were then used to parameterise simulations using the SL model, and the results analysed to provide the first direct estimates of *N_e_* reductions in natural *D. melanogaster* populations experiencing seasonal fluctuating selection.

### Estimating the genome-wide effect of seasonal fluctuating selection on N_e_ in D. melanogaster

To ensure that our simulations reproduce a feasible genomic context for *D. melanogaster*, we randomly introduced a specific number of seasonal loci reported in previous studies (ranging between 9 and 264 loci; Table 2) into one million diploid individuals each bearing the two main *D. melanogaster* autosomes (with empirically informed recombination rates from Comeron et al. (Comeron et al. 2012) and seasonal loci omitted from autosomal arm 3R to facilitate *N_e_* estimation at unlinked loci; see Methods). All simulated scenarios ran for 20,000 generations, with seasonal alleles introduced in the first generation at 50% frequency and neutral mutations introduced at the same frequency every 2,000 generations at 100kb intervals across both chromosomes (969 in total). For each sampled time point, we recorded neutral allele frequencies across all simulated generations over a complete seasonal cycle lasting 20 generations, and used the variation in neutral allele frequencies across the seasonal cycle to measure the instantaneous variance *N_e (Waples 1989; Jónás et al. 2016)_* (noting that generation and population size parameters were again downscaled by a factor of two to improve computational efficiency in the actual simulations, and neutral alleles were removed after each *N_e_* estimation). Finally, the impact of fluctuating selection on neutral diversity levels was inferred by evaluating *N_e_* relative to the census population size (i.e. *N_e_*/ *N_c_*).

Our simulations robustly reproduced the anticipated number of segregating seasonal alleles after 20,000 generations (all but one scenario falling within the 95% confidence interval, CI, of the expected empirical values; Table S1), though the mean seasonal frequency amplitude tended to be slightly lower than expected (reduced by 0.2-2% and only one scenario falling within the associated 95% CI), with the maximum discrepancy for both mean amplitude and final number of segregating loci occurring when simulating the most exaggerated seasonal fluctuations (i.e. 9 final loci with an average amplitude of 35%). These results suggest that our final *N_e_* estimates may be slightly conservative for each of the simulated scenarios.

The impact of fluctuating selection on the genome-wide *N_e_* was significant though highly variable, with reductions ranging between 7-76% relative to *N_c_* (Figure 2A & S7). These reductions reached stable values within the first 2,000 generations, concordant with the rapid stabilisation observed for frequency fluctuations. While the most pronounced *N_e_* reductions occurred prior to stable frequency oscillations being reached, this pattern likely represents an artefact of the highly skewed reproductive variances resulting from all seasonal alleles commencing at 50% frequencies.

**Figure 2.**
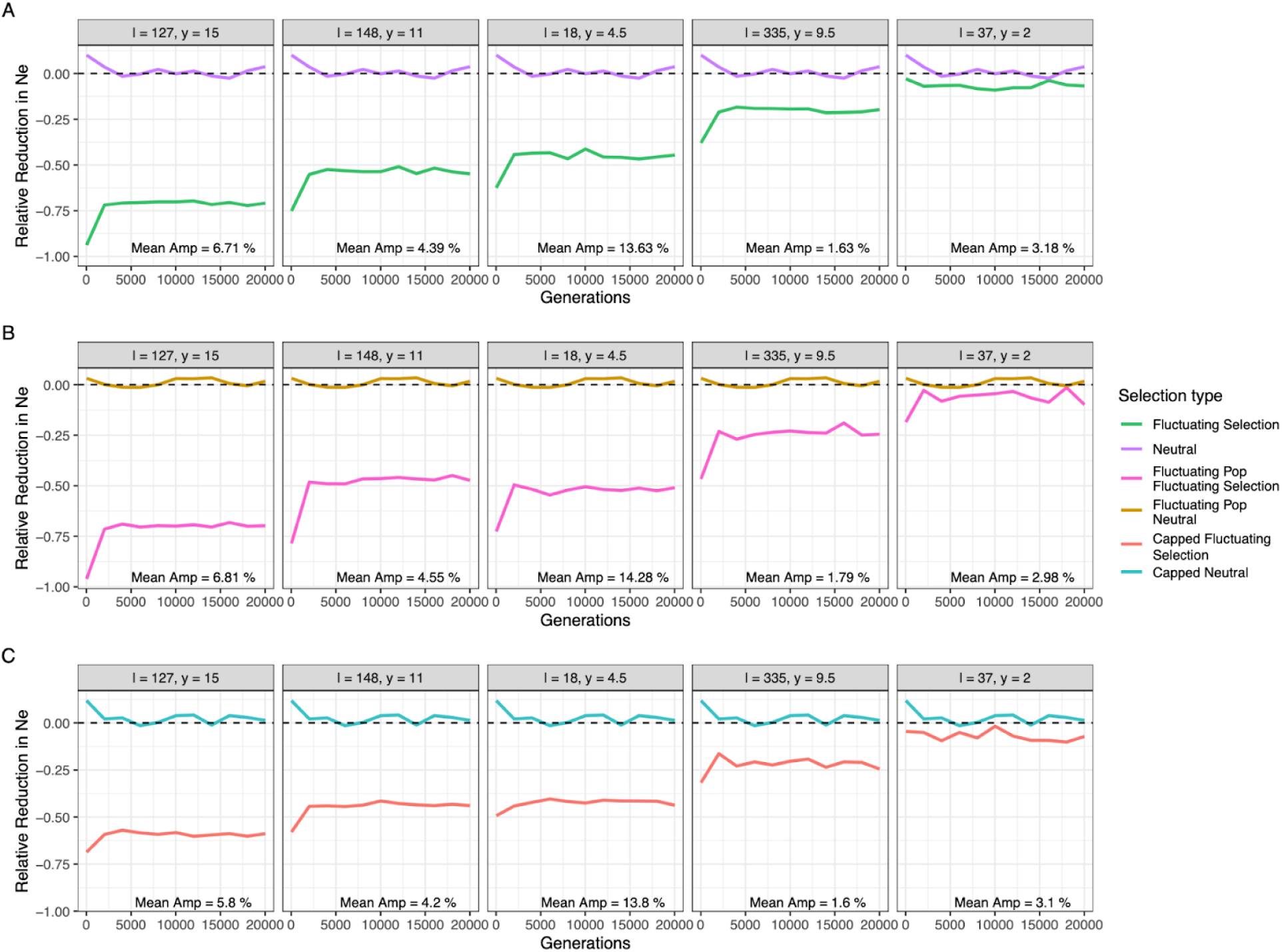
Relative reduction in effective population size due to fluctuating selection. Relative reduction in effective population size across time (generations) **A** under standard conditions with neutral evolution depicted in purple and fluctuating selection in green, **B** with seasonally fluctuating population size with gold and pink signifying neutrality and fluctuating selection, respectively, and **C** when offspring number was capped at 20 individuals with blue illustrating neutral evolution and fluctuating selection in red. Each panel shows effective population size for a different combination of initial loci number and epistasis value (labelled respectively in the top strip). A dashed black line depicts the neutral expectation (i.e. 0% reduction). The corresponding mean amplitude of each parameter combination is included in the bottom right of each panel.

The magnitude of *N_e_* reduction also depended on the sampling time within seasons. We calculated *N_e_* every generation over a seasonal cycle and observed the largest changes occurring following the seasonal transition due to increased reproductive variance over this period – though this effect is not readily discernible in scenarios where the *N_e_* reductions were low (Figure 3 & S8). Notably, *N_e_* reductions calculated across single generations within a seasonal cycle have reduced magnitudes compared to those calculated over larger timescales as *N_e_* estimated over a single generation does not adequately capture genetic drift occurring over a full seasonal cycle (Figure S8). A similar downward bias has been reported for variance-based estimators of *N_e_* when measurements are made over relatively few generations, seemingly due to the increased noise ratio in the allele frequency change estimates (Waples 2016, 2022).

**Figure 3.**
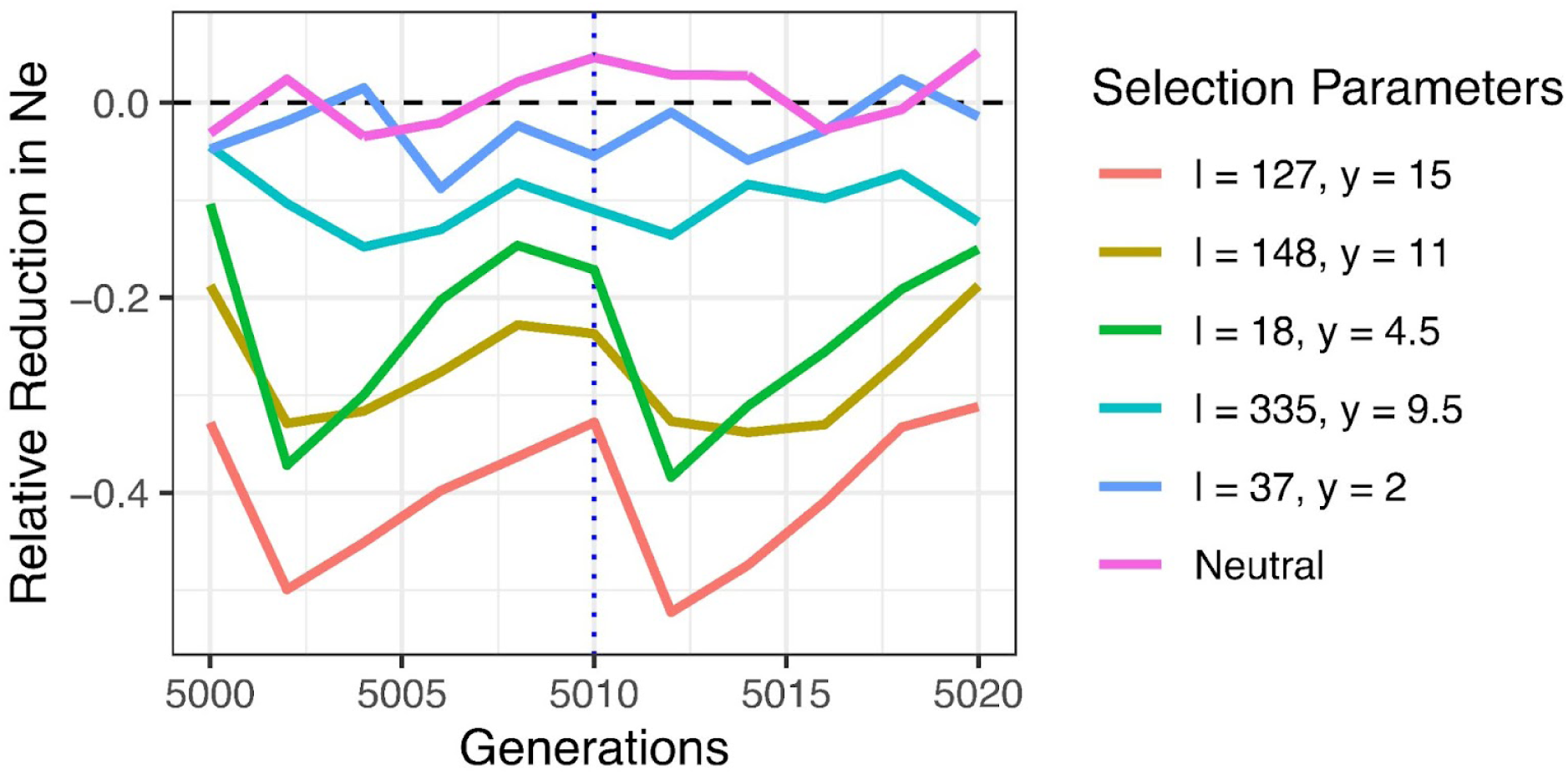
Effective population size across the seasonal cycle. Effective population size for different fluctuating selection parameters (coloured lines) across the seasonal cycle, 2 seasons consisting each of 10 generations. The blue, dashed vertical line signifies the time at which the seasons change. *N_e_* was sampled from generation 5000 to 5020 of the simulation as previous simulations suggested the reduction in effective population size had become stable at this time.

### Reductions in N_e_ scale with the largest seasonal allele frequency fluctuations

Interestingly, the greatest reduction in *N_e_* was not observed under scenarios with the most extreme mean seasonal fluctuations, but from the scenario where ∼90 seasonal loci stably oscillated with a mean amplitude of 6.7%, which closely aligns with the mean fluctuations reported in natural and outdoor experimental *Drosophila* populations (Machado et al. 2021; Rudman et al. 2022; Bitter et al. 2024). Further, while neither the mean nor the median seasonal amplitude showed a clear relationship with *N_e_* reduction – largely due to one scenario being an outlier (∼9 final loci, *y* = 4.5) – the maximum seasonal fluctuation displayed a strong linear correspondence with *N_e_* reduction (*r*^2^ = 0.702). This result suggests that seasonal loci with the most extreme allele frequency fluctuations are better predictors of *N_e_* reductions when fluctuating selection operates under a SL model.

Closer examination of the seasonal loci exhibiting the largest fluctuations showed a strong positive relationship with the mean effect size (Figure 4A) and little dependence on the mean dominance coefficient, which instead appeared to constrain the lower limits of the frequency amplitudes (Figure 4B). For all five of our modelled scenarios, the seasonal locus exhibiting the maximum frequency fluctuations tended to remain the same across time in each simulation (the only exceptions being a handful of simulations where the maximal amplitudes alternated between two loci). The close concordance between the maximum allele frequency fluctuation and *N_e_* prompted us to develop another linear regression model (equation 7; Methods) where expected *N_e_* reductions were calculated relative to the maximum seasonal frequency amplitudes. Applying this regression model to the maximum frequency amplitudes reported for natural *Drosophila* populations – i.e. 37% in Bergland and colleagues 2014 study and 45% in Machado et al’s follow up study in 2021 (Bergland et al. 2014; Machado et al. 2021) – resulted in predicted *N_e_* values that are ∼40-50% lower than the census population sizes (Figure 5). These results are consistent with the magnitude of reductions observed from our previous regression analyses and together suggest that the continual exposure of cosmopolitan *Drosophila* species to shifting seasonal selection pressures imposes a significant constraint on the effective size of the population that has been unappreciated previously.

**Figure 4.**
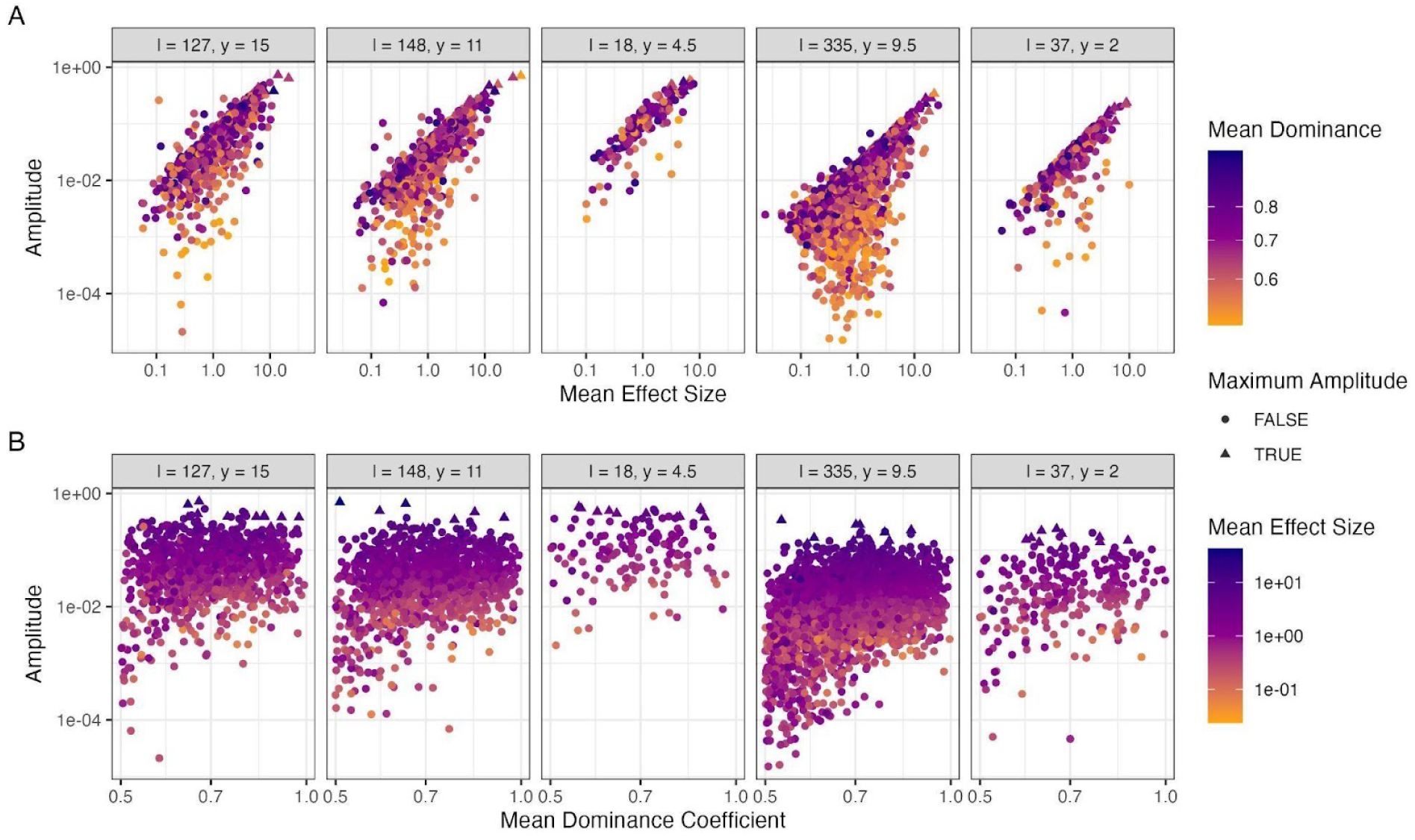
Mean effect size and mean dominance coefficient of seasonal loci segregating at the end of simulations in relation to their amplitude of fluctuation. **A** The mean effect size and **B** mean dominance coefficient, calculated as the average of the summer and winter value for each locus, compared to the amplitude of each locus. The colours of the points illustrate the mean dominance or effect size. Each panel depicts a different parameter combination labelled in the top strip with the initial loci number and epistasis value, respectively. Loci from all 10 simulated replicates are included in each panel. Loci identified as having the maximum amplitude at any point in a simulated replicate are depicted as triangles.

**Figure 5.**
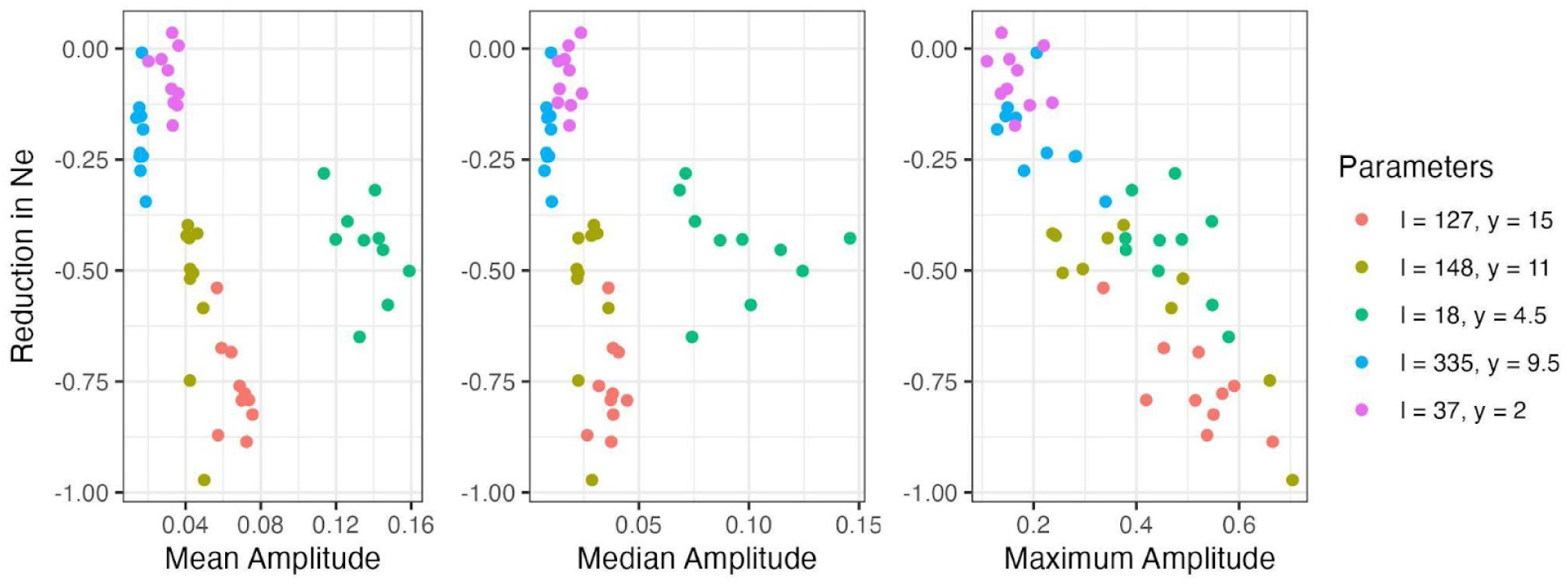
Relationship between reduction in effective population size and aspects of seasonal fluctuation (amplitude). The relative reduction in effective population size in relation to the mean, median, and maximum amplitude for each simulated replicate of each combination of initial loci number and epistasis value (coloured).

### Fluctuating selection produces consistently large genome-wide N_e_ reductions under alternate demographic scenarios

We explored the generality of our findings under different demographic models, first simulating fluctuating selection under a wide range of constant population sizes that spanned two orders of magnitude (i.e., *N_c_* ranging between 20 thousand to one million individuals) and replicating parameters from the scenario showing the greatest reduction in *N_e_*(75% for 127 initial loci and *y* = 15). Notably, the resulting *N_e_* reduction did not substantially change across all four examined *N_c_* values (Figure S9), indicating that the *N_e_* reduction is independent of *N_c_*.

Next, we allowed for population sizes to change by an order of magnitude between seasons to mimic the seasonal boom-bust cycles observed in *D. melanogaster* populations in temperate climates (i.e., we simulate sizes peaking at 1 million in summer and declining to 100,000 in winter). Rerunning each empirical parameter set with these alternating population sizes resulted in *N_e_* reductions that scaled with the harmonic mean population size, producing *N_e_* values an order of magnitude below the census population sizes in the summer period, or proportional decreases similar to the constant size model when scaled against the harmonic mean population size (Figure 2B).

Finally, previous theoretical work has shown that the SL model can produce unrealistically large offspring numbers under some scenarios, resulting in high reproductive variance where some individuals contribute unrealistically large offspring numbers to the subsequent generation (Wittmann et al. 2017). Accordingly, we reran our simulations imposing a limit on the maximum number of offspring per parent to 20, observing only a minor reduction in the simulated allele frequency fluctuations (Figure 2C) that agreed with the subtle change in the allele frequency fluctuations reported in theoretical results after imposing similar offspring caps (Wittmann et al. 2017). Further, this offspring cap had little effect on levels of *N_e_* reductions when the uncapped reduction was approximately 50% or less, though maximum reductions were limited to approximately 60% for scenarios where the uncapped *N_e_* reduction was more severe.

## Discussion

In recent years, fluctuating selection has emerged as a novel evolutionary factor that plays an important role in adapting cosmopolitan *Drosophila* populations and other species to their temporally varying habitats (Johnson et al. 2023). These findings have prompted the development of theoretical models to study the allele frequency dynamics of adaptively fluctuating loci and their broader impact on linked neutral variation (Barton 2000; Huerta-Sanchez et al. 2008; Taylor 2013; Wittmann et al. 2023, 2017). In this study, we show that fluctuating selection leads to *N_e_* values that are ∼50% smaller than expected under neutrality when simulating polygenetic seasonal selection in an empirically informed *Drosophila* model, with the scale of the decrease being primarily driven by the locus experiencing the largest frequency oscillations among all adaptively fluctuating loci. Our results clearly demonstrate the potent ability of fluctuating selection to reduce effective population size. This dramatic effect is a direct result of the exaggerated reproductive variance created by individuals repeatedly cycling between high and low fitness as environmental pressures continually alternate (Wittmann et al. 2023), as demonstrated by the greatest *N_e_* reductions occurring right after the transition to a new season when reproductive variance peaks (Figure 3).

Previous work investigating the effect of natural selection on effective population size has also included its consideration as a possible explanation for Lewontin’s Paradox (LP), the unexpected decoupling of neutral diversity and *N_c_* across populations (Lewontin 1974), due to its efficacy also increasing as population size increases (Lewontin 1974; Corbett-Detig et al. 2015; Roberts 2015; Buffalo 2021; Charlesworth & Jensen 2022; Achaz & Schertzer 2023; Kaushik 2023). While fluctuating selection has been proposed as a potential contributor to LP (Buffalo 2021), our simulations show that the resulting reductions in *N_e_* are largely independent of census size (*N_c_*). This indicates that fluctuating selection alone is insufficient to explain LP. This conclusion is consistent with recent work suggesting that the effects of linked selection, including recurrent selective sweeps, are generally too weak to account for the observed compression of genetic diversity across species (Coop 2016).

Nonetheless, fluctuating selection may still contribute to variation in the ratio of *N_e_* to *N_c_* among taxa. In particular, species such as arthropods, which experience strong and recurrent environmental fluctuations on generational timescales, may be especially affected due to the cumulative impact of temporally variable selection on reproductive variance. Unlike models based on recurrent positive selection, which require a sustained input of beneficial mutations and allow diversity to recover between sweeps (Buffalo 2021; Charlesworth & Jensen 2022; Achaz & Schertzer 2023), fluctuating selection can impose persistent reductions in *N_e_* as long as environmental conditions continue to alternate.

Together, these results suggest that fluctuating selection is unlikely to resolve LP on its own, but may act in concert with other processes, including demography and linked selection, to shape broad-scale patterns of genetic diversity.

### Conclusions and future work

Our findings emphasise that fluctuating selection is likely a fundamental, yet under-appreciated, determinant of effective population size among *Drosophila* and other species prone to temporal environmental shifts. Investigation of fluctuating selection is currently in its infancy, and our understanding of the breadth of fluctuating selection is hindered by the majority of prior empirical research focusing upon *Drosophila* species (Bergland et al. 2014; Machado et al. 2021; Behrman & Schmidt 2022; Rudman et al. 2022; Bitter et al. 2024; Nunez et al. 2024), despite other ectothermic species being obvious candidates for investigation due to their life-histories being highly sensitive to thermal environments (Pfenninger & Foucault 2022; Pfenninger et al. 2023; Lynch et al. 2024; Johnson et al. 2023). Beyond the need for more work on experimental and natural populations from diverse species, additional investigation is required to determine whether the *N_e_* effects observed in our simulations are robust to different models of fluctuating selection as these emerge in the future (Bertram & Masel 2019). Further, while our analyses show that *N_e_* reductions are relatively insensitive to changes in population size and restrictions on the number of offspring per parent, future research needs to examine the impact of other evolutionary and demographic forces, including seasonal asymmetries in generation number, continuous changes in population size, and the inclusion of background selection, as well as eco-evolutionary parameterisations tailored toward different species. This will enhance our understanding of how fluctuating selection shapes effective population size and contributes to genome-wide patterns of genetic variation across diverse species.

## Methods

### Parameter selection for the Segregation Lift model

The segregation lift model consists of two parts, a score of the contribution of the loci in a genotype (*z_s/w_*; see equation 1 and Table 1) and a fitness equation (see equation 2) (Wittmann et al. 2017). For the score component, each seasonal locus was allocated seasonal dominance coefficients (*d_s/w_*) and effect sizes (*Δ_s/w_*). Dominance coefficients for each season were drawn independently from a uniform distribution with a minimum of 0 and a maximum of 1, with only seasonal loci having mean dominance values greater 0.5 being retained because these loci are more likely to become stable polymorphisms (Wittmann et al. 2017). Effect sizes for each seasonal locus were drawn from a bivariate normal distribution with a mean of 0, standard deviation of 1, and correlation coefficient of 0.9, with both values being exponentiated to create a log-normal distribution (Wittmann et al. 2017). Correlated values were used since smaller differences between the summer and winter effect size are more likely to produce stable polymorphisms (Wittmann et al. 2017). The contribution of a locus was either the seasonal effect size, if homozygous for the seasonally favoured allele, the product of the seasonal effect size and the dominance coefficient if heterozygous, or 0 if homozygous for the unfavoured allele (Table 1). The seasonal score is calculated for each individual each generation as the sum of each loci’s contribution depending on the individual’s genotype (equation 1). This score is then used to calculate each individual’s fitness as per equation 2. Epistasis levels recreate a diminishing-returns fitness model (*y* range: 0.5, 1, 2, 4, 8, 12, 16, 20), as evidence for this model has appeared in several empirical studies (Chou et al. 2011; Khan et al. 2011; Kryazhimskiy et al. 2014) and theoretical studies show that it is more likely to produce stable fluctuating loci that experience reasonable reversals of dominance (whereas other forms of epistasis require almost complete reversal of dominance) (Wittmann et al. 2017). Finally, we followed empirical reports of the number of segregating seasonal loci in natural *Drosophila* populations (Rudman et al. 2022; Bitter et al. 2024) and introduced either 100, 200, or 500 seasonal loci (*l*) in our initial simulations. Importantly, our selection model assumes ‘soft’ selection, i.e. selection does not affect the census population size, and the *N_e_* reduction is solely attributable to the increased variance in offspring number.

**Table 1.** Contribution of seasonal loci (*C_l_)* to fitness.

| Season | $C_{l,ww}$ | $C_{l,sw}$ | $C_{l,ss}$ |
| --- | --- | --- | --- |
| Winter | $\Delta_w$ | $\Delta_w d_w$ | 0 |
| Summer | 0 | $\Delta_s d_s$ | $\Delta_s$ |

The contribution of a seasonal locus is conditioned on the genotype at that locus and the current season. If the locus is homozygous for the seasonal allele the contribution is the effect size of the seasonal allele. If the locus is homozygous for the allele of the opposing season the contribution is 0. If the locus is heterozygous, the contribution is the product of the effect size and the dominance coefficient for that given season.

### Forward simulations of fluctuating allele frequency trajectories

All simulations were performed using *SLiM* (v 4.0.1; (Haller & Messer 2023)), starting with simulations designed to quantify the allele frequency trajectories of unlinked seasonal loci under the SL model. Simulations were performed on different combinations of seasonal loci number and epistasis intensities (*y* range: 0.5, 1, 2, 4, 8, 12, 16, 20; *l* range: 100, 200, 500), using constant population sizes of 500,000 diploid individuals and 5 generations per season, and running for 18,000 generations in total. Note that this parameterisation produces allele frequency dynamics that are equivalent to 36,000 generations of evolution in a population of 1 million individuals experiencing 10 generations per season, which are broadly commensurate with reported parameters from natural *Drosophila* populations. Due to computational constraints, for simulations starting with 500 loci, we only simulated epistasis intensities of 8, 12 and 20, as preliminary simulations performed on smaller sized populations suggested that this range produced results most consistent with empirical data.

A second set of allele frequency simulations using reduced population sizes were also performed to provide increased resolution for the allele frequency dynamics of seasonal loci over the initial phase of selection. For these simulations, population size was downscaled by a factor of 100 (i.e. 10,000 individuals) to expedite computational requirements and run times, with all other parameter ranges remaining unchanged.

### Multiple linear regression of allele frequency amplitude for estimating selection parameters

We used the information obtained in the allele frequency simulations to develop a quantitative model of epistasis intensity (*y*) as a function of the number of seasonal loci and the amplitude of seasonal allele frequency change. We alternatively used the mean, median, and 90% quartile of seasonal loci frequency changes as one of the independent variables, with the mean amplitude giving the best model-fit overall (*r*^2^ = 0.988; *cf.* median: *r*^2^ = 0.963, 90% quartile: *r*^2^ = 0.985). An ANOVA model comparison suggested that adding a two-way interaction term significantly improved the model-fit (*p* < 0.001; *r*^2^ = 0.989; see equation 3, where *y* is the epistasis value, *n* is the final number of segregating seasonal loci, and *x* is the mean amplitude of seasonal allele frequency fluctuation) and this model was subsequently used to predict epistasis (*y*).

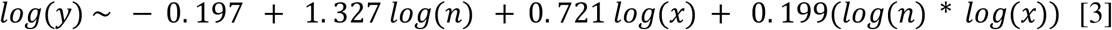

A multiple regression model was also used to determine the number of initially segregating loci required to obtain the final number of segregating loci after seasonal allele frequency oscillations had stabilised. This model predicts the required initial number of segregating loci (*l*) from the epistasis parameter (*y*) and final number of segregating loci (*n*). Again, adding an interaction term to the model (equation 4) significantly improved the fit (*p* < 0.001).

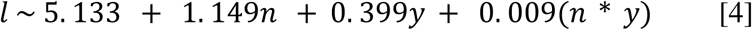

Using the two fitted regression models shown in equations 3 and 4, we predicted the epistasis intensities and the initial number of segregating loci needed to replicate the mean amplitude of allele frequency fluctuation and the number of segregating loci reported in studies of natural *D. melanogaster* populations (Table 2). For the final number of segregating seasonal loci, we took values reported in recently published outdoor *D. melanogaster* cage experiments (Rudman et al. 2022; Bitter et al. 2024), since the authors used linkage disequilibrium, hitchhiking, and parallel allele frequency changes to provide a conservative estimate of the number of independent seasonal loci coming under selection. For mean frequency amplitude, we used a range of values obtained from natural and experimental *D. melanogaster* studies (Bergland et al. 2014; Machado et al. 2021; Rudman et al. 2022; Bitter et al. 2024), utilising criteria that are summarised in Table 2.

**Table 2.**
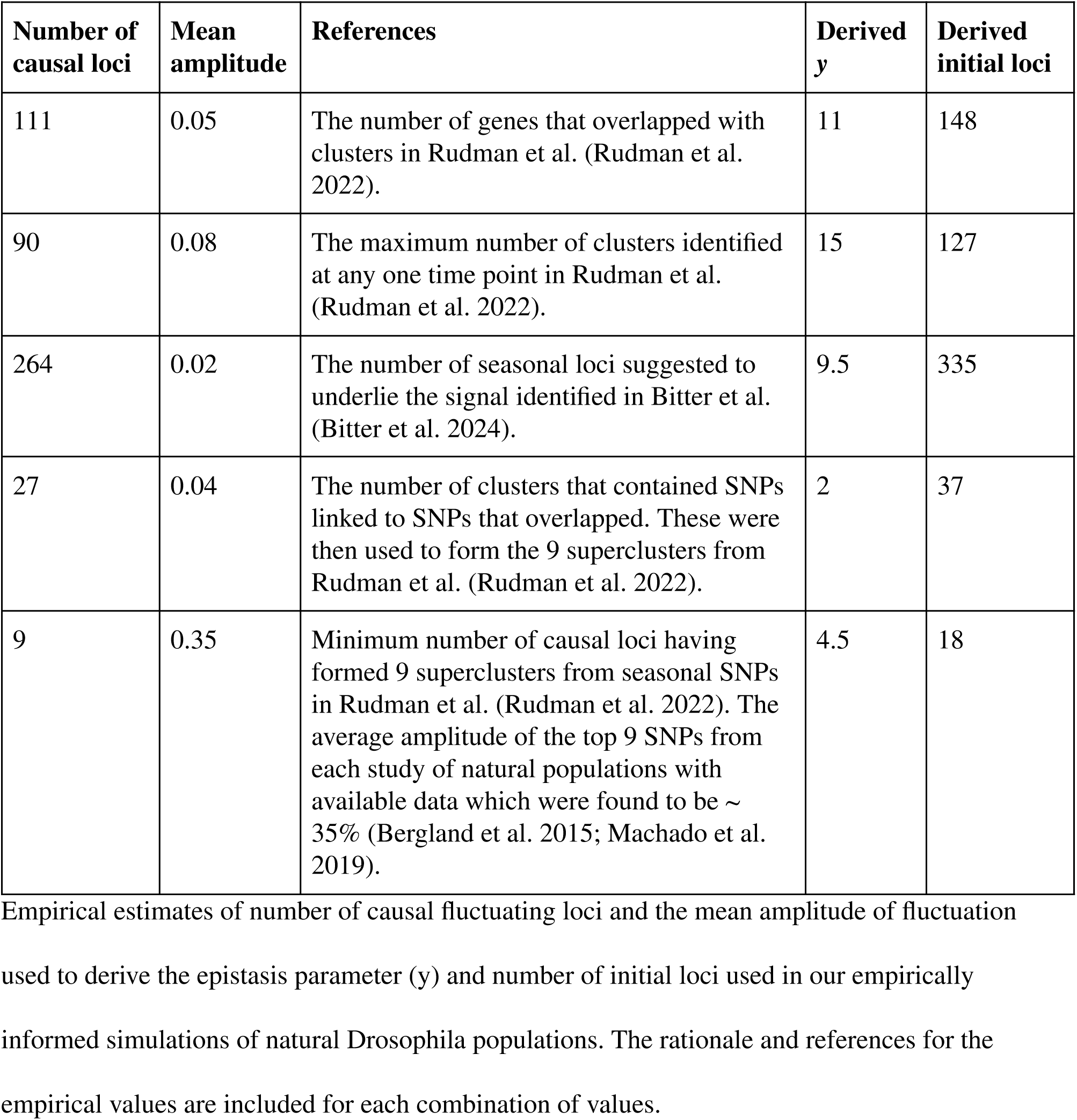
Parameters for genome-wide simulations.

### Empirically informed simulations of natural Drosophila populations

We used the empirically derived parameters estimated from our multiple linear regression models to simulate fluctuating selection under the SL model in diploid individuals that each carried both *Drosophila* autosomes, with chromosome lengths taken from the FlyBase (Jenkins et al. 2022) and recombination rates following Comeron et al. (Comeron et al. 2012). At the start of the simulations, seasonal alleles were introduced at 50% frequency at random positions across three of the four chromosome arms, leaving the final chromosomal arm (3R) with no seasonal loci to facilitate recording of neutral allele frequency changes on a genomic region unlinked to selected loci. Simulations used constant population sizes of 500,000 diploid individuals and over both seasons in each year, with 5 generations per season, and running for 10,000 generations in total, producing dynamics that are evolutionarily equivalent to 20,000 generations of evolution in a population of 1 million individuals experiencing 10 generations per season.

We also ran additional simulations that mimicked the boom-bust demography observed in cosmopolitan *Drosophila*, where the winter population size was reduced to 100,000 (i.e. 50,000 individuals after appropriate downscaling).

### Effective population size estimation

To measure the reduction in in *N_e_* caused by fluctuating selection in the empirically informed *Drosophila* simulations, neutral mutations starting at 50% frequency were inserted every 100 kb across the simulated genomes at multiple sampling timepoints, and the standardised variance, *F,* was calculated for each neutral mutation after *g* generations of change, according to equation 5 (Waples 1989).

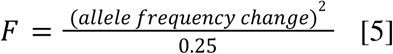

These *F* values were averaged separately across chromosome arms with and without seasonal loci (i.e., 2L, 2R, 3L or 3R) and used in equation 6 to calculate the *N_e_*, with *g* signifying the number of generations (i.e. 10 in this case) (Waples 1989; Jónás et al. 2016).

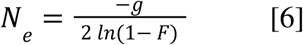

Neutral mutations were removed from the simulation following the estimation of *N_e_* and reintroduced at 50% at the next sampling point.

To capture *N_e_* changes across successive generations of the seasonal cycle, we ran a separate set of simulations for 2,500 generations (i.e., 5,000 unscaled), and we inserted neutral mutations starting at 50% frequency in each of the next 10 simulated generations (i.e. a single seasonal cycle, successively removing alleles from the previous generation) and measured *N_e_* using equation 6 (where *g* = 1). To examine if the discrepancy observed between *N_e_* calculated over 1 or 10 simulated generations was an artefact (Waples 2016, 2022), we repeated this process but let each neutral locus segregate for 30 generations and calculated *N_e_* at each generation (i.e., *g* = 1, 2, 3, …, 30), limiting calculations to loci that were still segregating after 3 complete seasonal cycles.

### Estimating reduction in N_e_ from maximum allele frequency amplitudes

We used standard linear regression to quantify the relationship between maximum allele frequency amplitude and the reduction in *N_e_*, obtaining the model shown in equation 7.

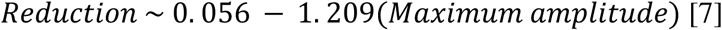

We used this equation to estimate the reduced *N_e_* in natural *Drosophila* populations previously reported to be under fluctuating selection (Bergland et al. 2015; Machado et al. 2019), obtaining maximum allele frequencies from data retrieved from the relevant Dryad repositories (code available at https://github.com/olivia-johnson/FluctuatingSelectionNe).

### Offspring Capping

Our simulation of strong seasonal selection can produce an unrealistically large number of offspring for a small number of highly fit parental individuals. We capped the number of offspring at 10 simulated individuals (i.e. 20 individuals prior to parameter downscaling), using *SLiM*’s ‘modifyChild’ callback. This allowed alternate parents to be allocated to individuals of a new generation if the ones originally picked had already contributed to 10 individuals.

## Author Contributions

C.D.H. led the study design and supervised the project. All authors contributed to the study design. O.L.J. set up and ran the simulations. O.L.J. conducted the majority of the analysis, with contributions from C.D.H. and R.T. All authors contributed to interpreting the results and writing the manuscript.

## Acknowledgements

Thanks go to Ben Haller for assistance with SLiM code and bug fixes; to Fabien Vosin, Gludhug Purnomo, Xavier Roca-Rada and Shyamsundar Ravishankar for their assistance with computational issues; and to Meike Wittmann and Yuseob Kim for their insightful comments on this manuscript.

## Data Availability Statement

The code to replicate the simulations, data processing, and analysis can be found at https://github.com/olivia-johnson/FluctuatingSelectionNe.

## References

Achaz G, Schertzer E. 2023. Weak genetic draft and the Lewontin’s paradox. bioRxiv. 2023.07.19.549703. doi: 10.1101/2023.07.19.549703.

Barton NH. 2000. Genetic hitchhiking. Phil. Trans. R. Soc. Lond. B. 355:1553–1562. doi: 10.1098/rstb.2000.0716.

Behrman EL et al. 2018. Rapid seasonal evolution in innate immunity of wild Drosophila melanogaster. Proc. Biol. Sci. 285:20172599. doi: 10.1098/rspb.2017.2599.

Behrman EL, Schmidt P. 2022. How predictable is rapid evolution? bioRxiv. 2022.10.27.514123. doi: 10.1101/2022.10.27.514123.

Bergland AO, Behrman EL, O’Brien KR, Schmidt PS, Petrov DA. 2015. Data from: Genomic evidence of rapid and stable adaptive oscillations over seasonal time scales in Drosophila. doi: 10.5061/DRYAD.V883P.

Bergland AO, Behrman EL, O’Brien KR, Schmidt PS, Petrov DA. 2014. Genomic evidence of rapid and stable adaptive oscillations over seasonal time scales in Drosophila. PLoS Genet. 10:e1004775. doi: 10.1371/journal.pgen.1004775.

Bertram J, Masel J. 2019. Different mechanisms drive the maintenance of polymorphism at loci subject to strong versus weak fluctuating selection. Evolution. 73:883–896. doi: 10.1111/evo.13719.

Bitter MC et al. 2024. Continuously fluctuating selection reveals fine granularity of adaptation. Nature. 634:389–396. doi: 10.1038/s41586-024-07834-x.

Buffalo V. 2021. Quantifying the relationship between genetic diversity and population size suggests natural selection cannot explain Lewontin’s paradox. eLife. 10:e67509. doi: 10.7554/eLife.67509.

Charlesworth B. 2012. The role of background selection in shaping patterns of molecular evolution and variation: evidence from variability on the Drosophila X chromosome. Genetics. 191:233–246. doi: 10.1534/genetics.111.138073.

Charlesworth B, Jensen JD. 2022. How can we resolve Lewontin’s Paradox? Genome Biol. Evol. 14:evac096. doi: 10.1093/gbe/evac096.

Charlesworth D. 2006. Balancing selection and its effects on sequences in nearby genome regions. PLoS Genet. 2:e64. doi: 10.1371/journal.pgen.0020064.

Chou H-H, Chiu H-C, Delaney NF, Segrè D, Marx CJ. 2011. Diminishing returns epistasis among beneficial mutations decelerates adaptation. Science. 332:1190–1192. doi: 10.1126/science.1203799.

Comeron JM. 2014. Background selection as baseline for nucleotide variation across the Drosophila genome. PLoS Genet. 10:e1004434. doi: 10.1371/journal.pgen.1004434.

Comeron JM. 2017. Background selection as null hypothesis in population genomics: insights and challenges from *Drosophila* studies. Philos. Trans. R. Soc. Lond. B Biol. Sci. 372:20160471. doi: 10.1098/rstb.2016.0471.

Comeron JM, Ratnappan R, Bailin S. 2012. The many landscapes of recombination in Drosophila melanogaster. PLoS Genet. 8:e1002905. doi: 10.1371/journal.pgen.1002905.

Coop G. 2016. Does linked selection explain the narrow range of genetic diversity across species? bioRxiv. 042598. doi: 10.1101/042598.

Corbett-Detig RB, Hartl DL, Sackton TB. 2015. Natural selection constrains neutral diversity across a wide range of species. PLoS Biol. 13:e1002112. doi: 10.1371/journal.pbio.1002112.

Crow JF. 1954. Breeding structure of populations. II. Effective population number. Statistics and mathematics in biology.

Dobzhansky T. 1947. Adaptive changes induced by natural selection in wild populations of *Drosophila*. Evolution. 1:1–16. doi: 10.1111/j.1558-5646.1947.tb02709.x.

Gillespie JH. 1991. The causes of molecular evolution. Oxford University Press.

Haldane JBS, Jayakar SD. 1963. Polymorphism due to selection of varying direction. J. Genet. 58:237–242. doi: 10.1007/BF02986143.

Haller BC, Messer PW. 2023. SLiM 4: Multispecies eco-evolutionary modeling. Am. Nat. 201:E127–E139. doi: 10.1086/723601.

Huang Y, Wright SI, Agrawal AF. 2014. Genome-wide patterns of genetic variation within and among alternative selective regimes. PLoS Genet. 10:e1004527. doi: 10.1371/journal.pgen.1004527.

Huerta-Sanchez E, Durrett R, Bustamante CD. 2008. Population genetics of polymorphism and divergence under fluctuating selection. Genetics. 178:325–337. doi: 10.1534/genetics.107.073361.

Jenkins VK, Larkin A, Thurmond J, FlyBase Consortium. 2022. Using FlyBase: A database of Drosophila genes and genetics. Methods Mol. Biol. 2540:1–34. doi: 10.1007/978-1-0716-2541-5_1.

Johnson OL, Tobler R, Schmidt JM, Huber CD. 2023. Fluctuating selection and the determinants of genetic variation. Trends Genet. 39:491–504. doi: 10.1016/j.tig.2023.02.004.

Jónás Á, Taus T, Kosiol C, Schlötterer C, Futschik A. 2016. Estimating the effective population size from temporal allele frequency changes in experimental evolution. Genetics. 204:723–735. doi: 10.1534/genetics.116.191197.

Kaushik S. 2023. Effect of beneficial sweeps and background selection on genetic diversity in changing environments. J. Theor. Biol. 562:111431. doi: 10.1016/j.jtbi.2023.111431.

Kelly JK. 2022. The genomic scale of fluctuating selection in a natural plant population. Evol Lett. 6:506–521. doi: 10.1002/evl3.308.

Khan AI, Dinh DM, Schneider D, Lenski RE, Cooper TF. 2011. Negative epistasis between beneficial mutations in an evolving bacterial population. Science. 332:1193–1196. doi: 10.1126/science.1203801.

Kryazhimskiy S, Rice DP, Jerison ER, Desai MM. 2014. Microbial evolution. Global epistasis makes adaptation predictable despite sequence-level stochasticity. Science. 344:1519–1522. doi: 10.1126/science.1250939.

Lewontin RC. 1974. The genetic basis of evolutionary change. Columbia University Press https://play.google.com/store/books/details?id=rLMTAQAAIAAJ.

Lynch M, Wei W, Ye Z, Pfrender M. 2024. The genome-wide signature of short-term temporal selection. Proc. Natl. Acad. Sci. U. S. A. 121:e2307107121. doi: 10.1073/pnas.2307107121.

Machado HE et al. 2021. Broad geographic sampling reveals the shared basis and environmental correlates of seasonal adaptation in Drosophila. eLife. 10:e67577. doi: 10.7554/eLife.67577.

Machado HE et al. 2019. Data from: Broad geographic sampling reveals predictable, pervasive, and strong seasonal adaptation in Drosophila. doi: 10.5061/DRYAD.4R7B826.

Nunez JCB et al. 2024. A cosmopolitan inversion facilitates seasonal adaptation in overwintering Drosophila. Genetics. 226:iyad207. doi: 10.1093/genetics/iyad207.

Pfenninger M, Foucault Q. 2022. Population genomic time series data of a natural population suggests adaptive tracking of fluctuating environmental changes. Integr. Comp. Biol. 62:1812–1826. doi: 10.1093/icb/icac098.

Pfenninger M, Foucault Q, Waldvogel A-M, Feldmeyer B. 2023. Selective effects of a short transient environmental fluctuation on a natural population. Mol. Ecol. 32:335–349. doi: 10.1111/mec.16748.

Roberts RG. 2015. Lewontin’s paradox resolved? In larger populations, stronger selection erases more diversity. PLoS Biol. 13:e1002113. doi: 10.1371/journal.pbio.1002113.

Rodríguez de Cara MÁ et al. 2023. Balancing selection at a wing pattern locus is associated with major shifts in genome-wide patterns of diversity and gene flow. Peer Community J. 3. doi: 10.24072/pcjournal.298.

Rudman SM et al. 2022. Direct observation of adaptive tracking on ecological time scales in *Drosophila*. Science. 375:eabj7484. doi: 10.1126/science.abj7484.

Sjödin P, Kaj I, Krone S, Lascoux M, Nordborg M. 2005. On the meaning and existence of an effective population size. Genetics. 169:1061–1070. doi: 10.1534/genetics.104.026799.

Taylor JE. 2013. The effect of fluctuating selection on the genealogy at a linked site. Theor. Popul. Biol. 87:34–50. doi: 10.1016/j.tpb.2013.03.004.

Wakeley J, Sargsyan O. 2009. Extensions of the coalescent effective population size. Genetics. 181:341–345. doi: 10.1534/genetics.108.092460.

Wang J, Santiago E, Caballero A. 2016. Prediction and estimation of effective population size. Heredity. 117:193–206. doi: 10.1038/hdy.2016.43.

Waples RS. 1989. A generalized approach for estimating effective population size from temporal changes in allele frequency. Genetics. 121:379–391. doi: 10.1093/genetics/121.2.379.

Waples RS. 2016. Making sense of genetic estimates of effective population size. Mol. Ecol. 25:4689–4691. doi: 10.1111/mec.13814.

Waples RS. 2022. What is Ne, anyway? J. Hered. 113:371–379. doi: 10.1093/jhered/esac023.

Wittmann MJ, Bergland AO, Feldman MW, Schmidt PS, Petrov DA. 2017. Seasonally fluctuating selection can maintain polymorphism at many loci via segregation lift. Proc. Natl. Acad. Sci. U.S.A. 114:E9932–E9941. doi: 10.1073/pnas.1702994114.

Wittmann MJ, Mousset S, Hermisson J. 2023. Modeling the genetic footprint of fluctuating balancing selection: From the local to the genomic scale. Genetics. 223:iyad022. doi: 10.1093/genetics/iyad022.

